# Prefrontal cortex creates novel navigation sequences from hippocampal place-cell replay with spatial reward propagation

**DOI:** 10.1101/466920

**Authors:** Nicolas Cazin, Martin Llofriu Alonso, Pablo Scleidorovich Chiodi, Tatiana Pelc, Bruce Harland, Alfredo Weitzenfeld, Jean-Marc Fellous, Peter Ford Dominey

**Author notes:** **Corresponding Author**: Peter Ford Dominey.

## Abstract

As rats learn to search for multiple sources of food or water in a complex environment, they generate increasingly efficient trajectories between reward sites, across multiple trials. This optimization capacity has been characterized in the Traveling Salesrat Problem (TSP) (<u>de Jong et al (2011)</u>. Such spatial navigation capacity involves the replay of hippocampal place-cells during awake states, generating small sequences of spatially related place-cell activity that we call “snippets”. These snippets occur primarily during sharp-wave-ripple (SWR) events. Here we focus on the role of replay during the awake state, as the animal is learning across multiple trials. We hypothesize that snippet replay generates synthetic data that can substantially expand and restructure the experience available to make PFC learning more optimal. We developed a model of snippet generation that is modulated by reward, propagated in the forward and reverse directions. This implements a form of spatial credit assignment for reinforcement learning. We use a biologically motivated computational framework known as ‘reservoir computing’ to model PFC in sequence learning, in which large pools of prewired neural elements process information dynamically through reverberations. This PFC model is ideal to consolidate snippets into larger spatial sequences that may be later recalled by subsets of the original sequences. Our simulation experiments provide neurophysiological explanations for two pertinent observations related to navigation. Reward modulation allows the system to reject non-optimal segments of experienced trajectories, and reverse replay allows the system to “learn” trajectories that is has not physically experienced, both of which significantly contribute to the TSP behavior.

**Author Summary:** As rats search for multiple sources of food in a complex environment, they generate increasingly efficient trajectories between reward sites, across multiple trials, characterized in the Traveling Salesrat Problem (TSP). This likely involves the coordinated replay of place-cell “snippets” between successive trials. We hypothesize that “snippets” can be used by the prefrontal cortex (PFC) to implement a form of reward-modulated reinforcement learning. Our simulation experiments provide neurophysiological explanations for two pertinent observations related to navigation. Reward modulation allows the system to reject non-optimal segments of experienced trajectories, and reverse replay allows the system to “learn” trajectories that it has not physically experienced, both of which significantly contribute to the TSP behavior.

## Introduction

Spatial navigation in the rat involves the replay of place-cell subsequences (snippets) during awake and sleep states in the hippocampus during sharp-wave-ripple (SWR) events (<u>Carr et al 2011</u>, <u>Davidson et al 2009</u>, <u>Euston et al 2007</u>, <u>Kudrimoti et al 1999</u>). We focus on the role of replay during the awake state (Karlsson & Frank 2009), as the animal generates increasingly efficient trajectories between reward sites, across multiple trials. This trend toward near-optimal solutions is reminiscent of the classic Traveling Salesperson Problem (TSP) (<u>de Jong et al 2011</u>). While it appears likely that replay contributes to this learning behavior, the underlying neurophysiological mechanisms remain to be understood.

One obvious advantage of replay would be to provide extra training examples to otherwise slow reinforcement learning systems. This approach has been previously exploited with good results (<u>Johnson & Redish 2005</u>). We will go beyond this by prioritizing replay based on a spatial gradient of reward proximity that is built up during replay. We hypothesize (a) that snippet replay allows recurrent dynamics in prefrontal cortex (PFC) to consolidate snippet representations into novel efficient sequences, by rejecting other sequences that are less robustly coded in the input, and (b) that a form of reward-modulated replay in hippocampus implements a simple and efficient form of reinforcement learning to achieve this (Singer & Frank 2009).

An example of the behavior in question is illustrated in Figure 1. Panel A illustrates the optimal path linking the 5 feeders (ABCDE) in red. Panels B-D illustrate navigation trajectories that contain subsequences of the optimal path (in red), as well as non-optimal subsequences (in blue). In the framework of reward modulated replay, snippets from the efficient subsequences in panels B-D will be replayed more frequently, and will lead the system to autonomously generate the optimal sequence as illustrated in panel A. We thus require a sequence learning system that can re-assemble the target sequences from these replayed snippets. For this we choose a biologically inspired recurrent network model of prefrontal cortex (<u>Dominey 1995</u>, <u>Enel et al 2016</u>) that we believe will be able to integrate snippets from examples of non-optimal trajectories and to synthesize an optimal path.

**Figure 1:**
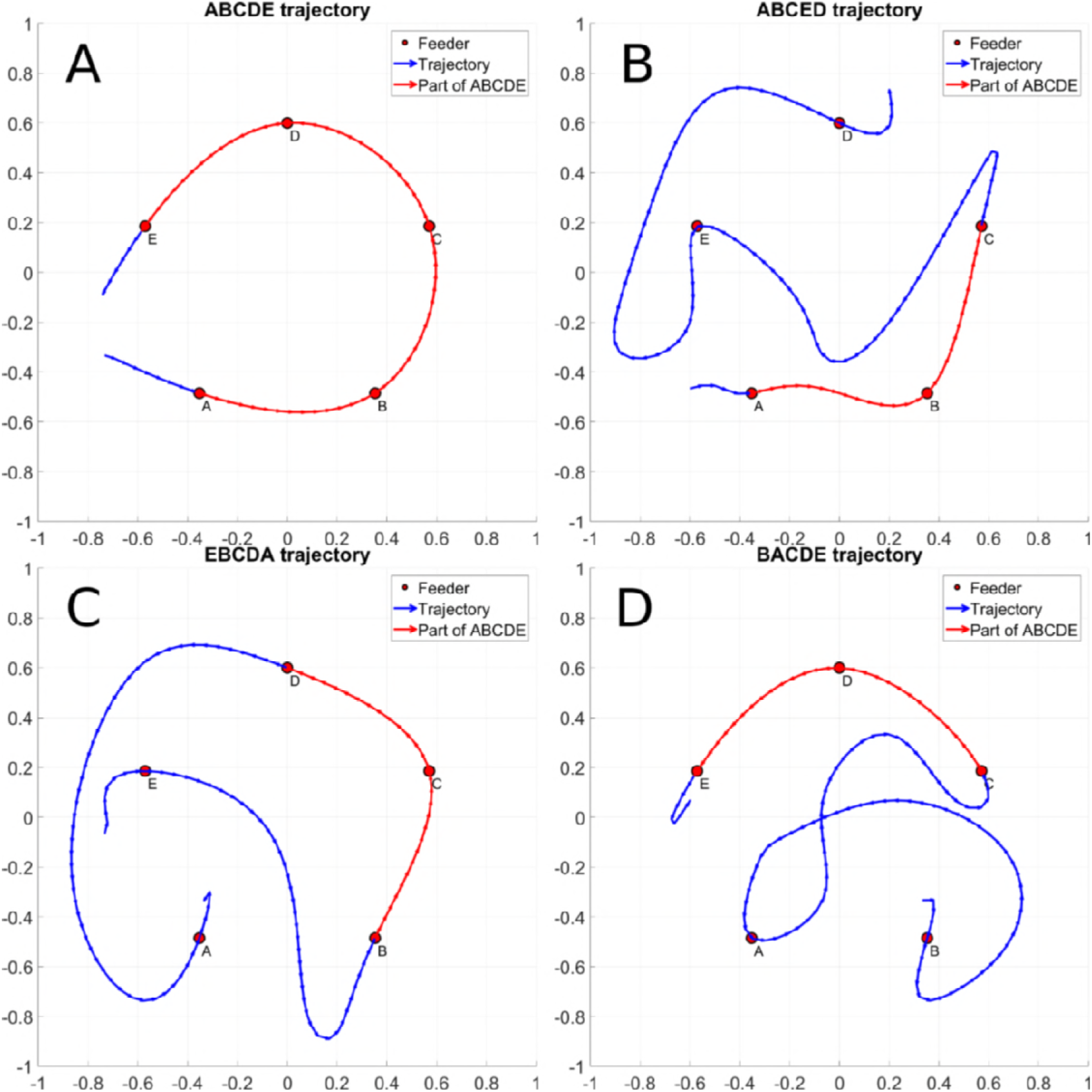
An optimal trajectory between feeders <u>ABCDE</u> is represented in panel A. Panel B, C and D display non optimal trajectories that contain a sub trajectory of the <u>ABCDE</u> trajectory. The sub trajectory shared with the <u>ABCDE</u> trajectory is displayed in red and the non-optimal parts in blue. Panel B contains the <u>ABC</u>ED, panel C the <u>EBCDA</u> trajectory and panel D the BA<u>CDE</u> trajectory.

For sequence learning, recurrent networks provide inherent sensitivity to serial and temporal structure. Modification of recurrent connections requires different methods of unwinding the recurrent connections in time, which limits the full dynamics of the recurrent system over extended time. To avoid this temporal cut-off and the space and time complexity required in the calculation of credit assignment to recurrent connections we used the framework of reservoir computing in which input and recurrent connections are fixed, and learning-related plasticity occur outside the reservoir network (Dominey 1995). The readout connections from the recurrent network learn the statistical structure of the data that the system is trained on, which places requirements on the mechanism that trains the model. We test the hypothesis that the structure of snippet replay from the hippocampus will provide the PFC with constraints that can be integrated in order to contribute to solving the TSP problem.

Two principal physical and neurophysiological properties of navigation and replay are exploited by the model and contribute to the system’s ability to converge onto an acceptable solution to the TSP. First, during navigation between baited food wells in the TSP task, non-optimal trajectories by definition cover more distance between rewards than near-optimal ones. Second, during the replay of recently activated places cells, the trajectories are encoded in forward and reverse directions (<u>Diba & Buzsaki 2007</u>, <u>Foster & Wilson 2006</u>). Exploiting these observations, we test the hypotheses that:

1. With replay biased by distance to reward, non-optimal trajectories will be less represented in replay, allowing the PFC to eliminate non-optimal subsequences in constructing the final efficient trajectory.
2. Reverse replay will allow the model to exploit the information provided by a given sequence in forward and backward directions, whereas the actual trajectory run by the rat has one direction only.

In testing these hypotheses, we will illustrate how the system can meet the following challenges:

1. Learn a global place-cell activation sequence from an unordered set of snippets
2. Consolidate multiple non-optimal sequences into a trajectory that efficiently links rewarded locations, thus converging to a good solution to the TSP problem.
3. Experience a trajectory in the forward direction and the learn to generate it in forward and backward direction, including concatenating parts of both forward and reverse replayed snippets in order to generate novel trajectories as demonstrated in <u>Gupta et al (2010)</u>.

The objective is to provide a coherent explanation of how critical aspects of replay – notably its modulation by reward, and the forward and reverse aspects, can be exploited by a cortical sequence learning system in order to display novel and efficient navigation trajectory generation.

The model developed in this research provides a possible explanation of mechanisms that allow PFC and hippocampus to interact to perform path optimization. This implies functional connectivity between these two structures. In a recent review of hippocampal–prefrontal interactions in memory-guided behavior <u>Shin and Jadhav (2016)</u> outlined a diverse set of direct and indirect connections that allow bi-directional interaction between these structures. Principal direct connections to PFC originate in the ventral and intermediate CA1 regions of the hippocampus (<u>Cenquizca & Swanson 2007</u>, <u>Harland et al 2018</u>). direct connections between hippocampus and PFC pass via the medial temporal lobe (subiculus, entorhinal cortex, peri- and post-rhinal cortex) (<u>Delatour & Witter 2002</u>), and the nucleus reuniens (<u>Vertes et al 2007</u>). Indeed memory replay is observed to be coordinated across hippocampus and multiple cortical areas including V1 (<u>Ji & Wilson 2007</u>). These studies allow us to consider that there are anatomical pathways that justify the modeling of bi-directional interaction between PFC and hippocampus (<u>McClelland et al 1995</u>).

## Material & methods

Experiments are performed on navigation trajectories (observed from rat behavior, or generated automatically) that represent the recent experience from the simulated rat. Snippets are extracted from this experience, and used to train the output connections of the PFC reservoir. This requires the specification of a model of place-cell activation in order to generate snippets. Based on this training, the sequence generation performance is evaluated to test the hypotheses specified. The evaluation requires a method for comparing sequences generated with expected sequences that is based on the Fréchet distance.

### Navigation Behavior and Trajectories

A trajectory is a sequence of N contiguous two-dimensional coordinates sampled from time *t*_1_ to time *t_N_* noted *L*(*t*_1_ → *t_N_*) that corresponds to the rat’s traversal of the baited feeders. The spatial resolution of trajectories depicted in is 20 *points/m* along the trajectory. Experiments were performed using navigation trajectories, including those displayed in Figure 1, based on data recorded from rats as they ran the TSP task (<u>de Jong et al 2011</u>) in a circular arena having a radius of 151cm. Twenty-one fixed feeders are distributed according to a spiral shape. In a typical configuration, 5 feeders are baited with a food pellet. For a given configuration, the rat runs several trials which are initially random and inefficient, and become increasingly efficient over successive trials, characterizing the TSP behavior (<u>de Jong et al 2011</u>). Rat data that characterizes the TSP behavior is detailed in S1, section Rat navigation data. The principle concept is that TSP behavior can be characterized as illustrated in Figure 1, where a system that is exposed to trajectories that contain elements of the efficient path can extract and concatenate these subsequences in order to generate the efficient trajectory.

### Place-cells

The modeled rat navigates in a closed space of 2×2 meters where it can move freely in all direction within a limited range (± 110° left and right of straight ahead), and encodes locations using hippocampus place-cell activity. A given location *s* = (*x*, *y*) is associated with a place-cell activation pattern by a set of 2D Gaussian place-fields:

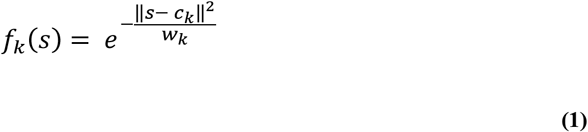

Where:

- *k* is the index of the place-cell
- *f_k_*(*s*) is the mean firing rate of the *k^th^* place-cell
- *c_k_* is the (*x*, *y*) coordinate of the *k^th^*place-cell
- 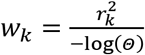 is a constant that will constrain the highest activations of the place-cell to be mostly contained in a circle of radius *r_k_*, centered in *c_k_*
- *r_k_* is the radius of the *k^th^* place-field
- *Θ* is the radius threshold which controls the spatial selectivity of the place-cell

Parameter *w_k_* is a manner of defining the variance of the 2D Gaussian surface with a distance to center related parameter *r_k_*. We model a uniform grid of 16×16 Gaussian place-fields of equal size (mimicking dorsal hippocampus). In Figure 2 the spatial position and extent of the place fields of several place-cells is represented in panel A by red circles. The degree of red transparency represents the mean firing rate.

**Figure 2:**
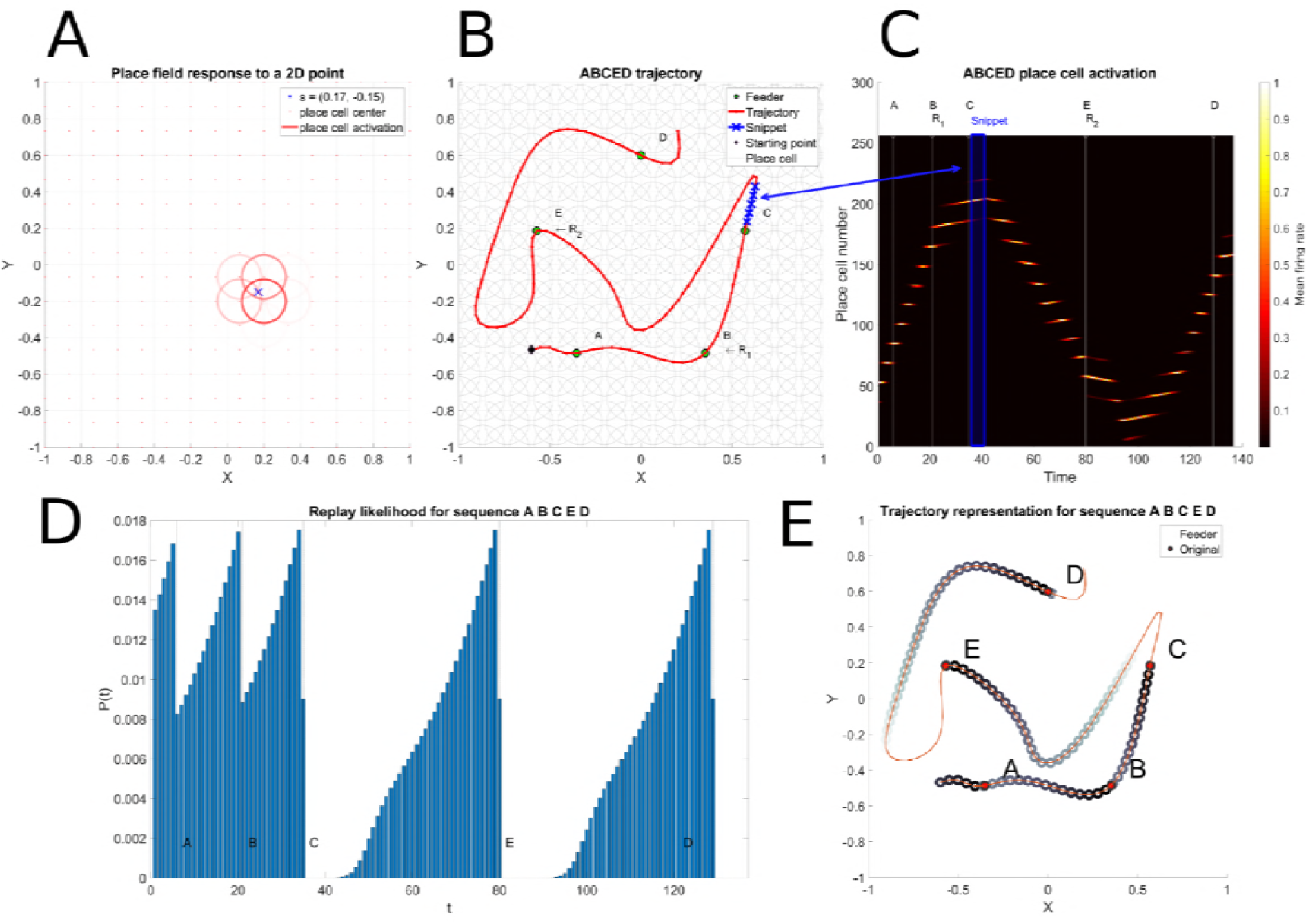
Place-cell and Snippet coding. Panel A represents the place-cell activations that correspond to a single point. Place-cell centers are represented by red points and the mean firing rate of each place-cell by a red circle with a fixed radius, centered on the place-cell center. The transparency level of the circle represents the magnitude of the mean firing rate. Panel B depicts the ABCED trajectory, the two randomly selected reward sites *R_1_* and *R_2_* and a snippet randomly drawn between *R_1_* and *R_2_*. The snippet length is *s = 5*. Panel C represents the raster of the place-cell activation along the ABCED trajectory. The time index where feeders A,B (*R_1_*), C,D and E(*R_2_*) are encountered during the ABCED trajectory are tagged above the raster and represented by a thin white vertical line. The snippet represented in panel B is emphasized by a blue rectangle in panel C. Panel D represents the snippet replay likelihood as learnt by the Hippocampal replay model and panel E represents the spatial extent of the snippet replay likelihood

A mean firing rate close to one will result in an bright circle if the location *s* is close to the place-field center *c_k_* of the place-cell *k*. For a more distant place-field center *c_l_* of place-cell *l*, the mean firing rate will be less important and the red circle representing this mean firing rate will be dimmer.

Thus at each time step, the place-cell coding that corresponds to a particular point in a trajectory is defined as the projection of this *L*(*t_n_*) point through *K* radial basis functions (i.e. Gaussian place-fields spatial response)

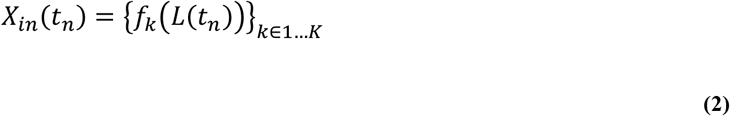

Each coordinate of the input vector *X_in_*(*t_n_*) represents the mean firing rate of hippocampus place-cells and its value lies between 0 and 1. Figure 2 represents in panel B the ABCED trajectory *L*(*t*_1_ → *t_N_*) and the corresponding place-cell mean firing rate raster *X_in_*(*t*_1_ → *t_N_*) is depicted in panel C

### Hippocampus replay

The hippocampus replay observed during SWR complexes in the active rest phase (between two trials in a given configuration of baited food wells) is modeled by generating condensed (time compressed) subsequences of place-cell activation patterns (snippets) that are then replayed at random so as to constitute a training set. The sampling distribution for drawing a random place-cell activation pattern might be uniform or modulated by new or rewarding experience as described in (<u>Carr et al 2011</u>). <u>Ambrose et al (2016)</u> show that during SWR sequences place-cell activation occur in reverse order at the end of a run. In particular, we model a random replay that is biased by reward. We will demonstrate an innovative method for spatial propagation of reward during replay that yields a computationally simple form of reinforcement learning.

We define a snippet as the concatenation of a pattern of successive place-cell activation:

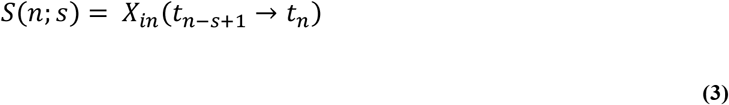

Where:

- *s* is the number of place-cell activations.

We define a time budget noted *T* that corresponds to the duration of a replay episode (experimentally, typically 70-100 ms). A replay episode *E* is a set of snippets of length *s*:

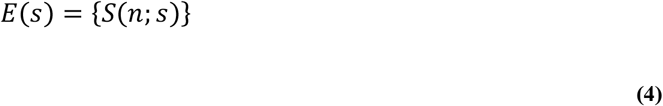

The sum of the durations of snippets replayed in *E* is constrained by *T*. If the time budget is exceeded one snippet is truncated in order to fit the time budget.

In Figure 2, Panel B represents a particular trajectory through feeders A, B, C, E and D. The depicted snippet is a subsequence of 5 contiguous locations belonging to the ABCED sequence. The B and E feeders are baited and marked as rewarding (R_1_ and R_2_). Panel B shows the spatial extent of a given snippet chosen in sequence ABCED and panel C shows the place-cell activation pattern of the ABCED trajectory and the corresponding snippet location in the raster.

#### Reward Propagation

The snippet replay model favors snippets that are on efficient paths linking rewarded sites (e.g. paths linking feeders A, B and C in panel B Figure 2), and not those that are on inefficient paths (as in paths linking C, D and E in the same panel). This is achieved by propagating reward value backwards from rewarded locations, and calculating the probability of replay as a function of proximity to a reward. Panel D of Figure 3 illustrates the resulting probability distributions for snippet selection along the complete path. Panel E represents the spatial extent of snippet replay likelihood. Note that the paths linking A, B and C have the highest probabilities for snippet replay.

**Figure 3:**
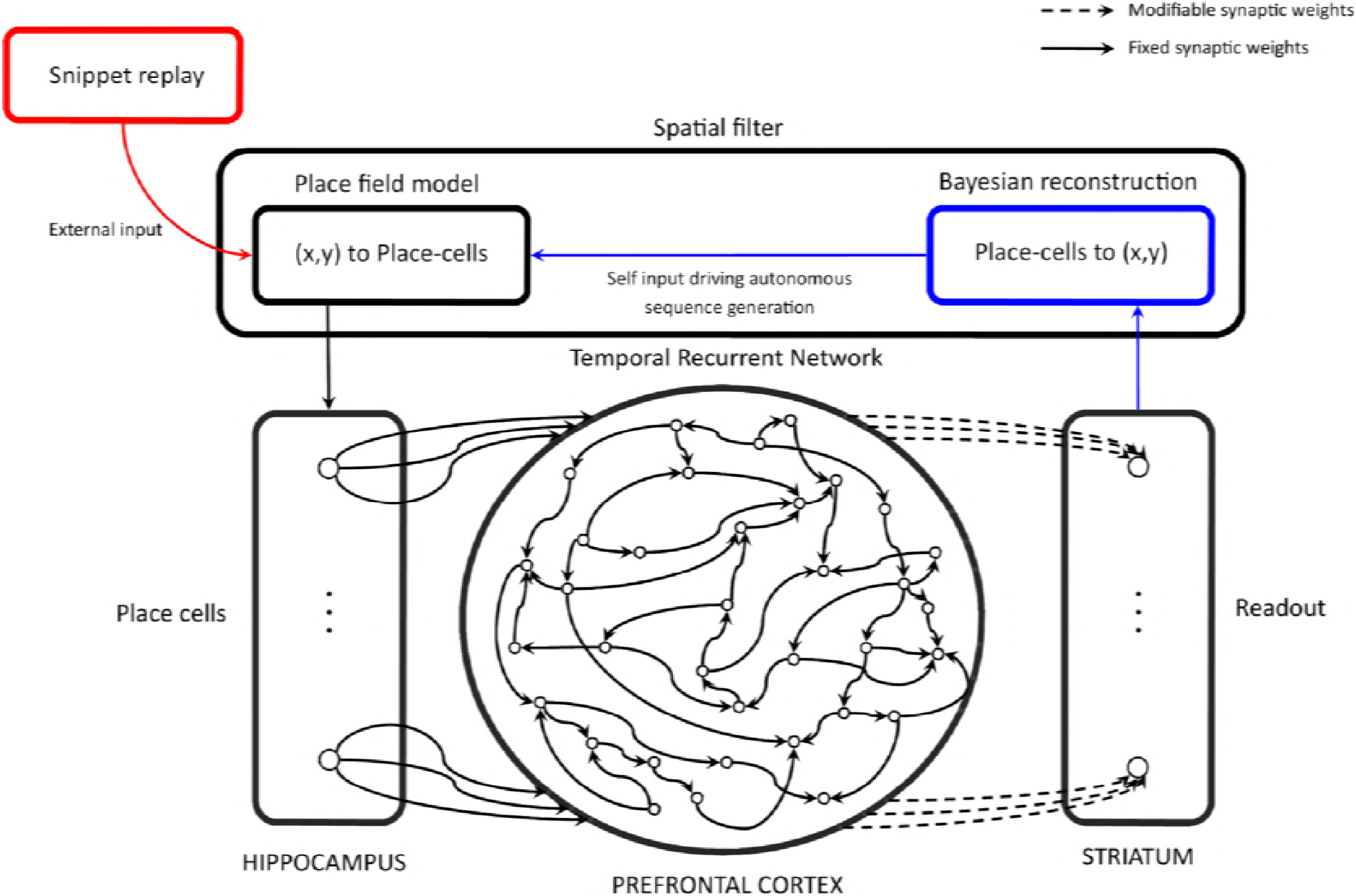
Reservoir Computing Model. The Temporal Recurrent Network (TRN) is a model of the prefrontal cortex (PFC) that take into account cortico-cortical loops by defining a fixed recurrent adjacency matrix for the leaky integrator neurons that model PFC neurons. Inputs of the TRN are modelled hippocampus (HIPP) place-cells. During the training phase, place-cells activations are provided by the algorithmic model of SWR replay (red pathway), and the striatum model learns to predict the next place-cell activation from the PFC model states by modifying the synaptic weights that project the PFC model into the striatum model according the delta learning rule. During the generation phase, the model is no longer learning and the place-cell activation patterns result from the new position of the agent, reconstructed with a Bayesian algorithm from the next place-cell activation prediction of the modeled striatum (blue pathway)

Hippocampus place-cell replay can occur in forward or backward direction as suggested in (<u>Foster & Wilson 2006</u>). We model the reverse replay as follows: For a given trajectory k of *N_k_* samples, there are *N_k_* − *s* possible snippets that can be replayed but only a limited number of snippets will be selected to fit the time budget *T*. A snippet *S*(*n*) has a likelihood of being replayed if it is related to a reward prediction. A generative model of snippet replay likelihood is first learnt by propagating a time delayed reward information according to the replay direction and the snippet duration. The timespan of a snippet acts as a propagation vector during the estimation phase of the snippet replay likelihood.

The reward prediction *v*(*t*_1_−> 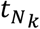) is learnt by initializing it to small positive random values and then iteratively refined by applying the procedure (8) K times:

1. Draw a random contiguous time index subset *τ* ≡ *T*(*n*, *s*, *r*; *β_learn_*) according to the reverse rate *β_learn_*:

a. Select a time-step *t_n_* such that *n* ∈ {1 … *N_k_*}
b. according to the replay likelihood defined by:

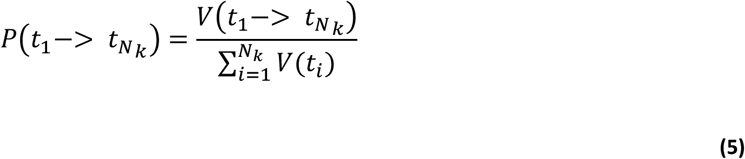
c. Select a random number *r* ∈ [0, 1] and a contiguous and monotonous time index sequence *τ* such that:

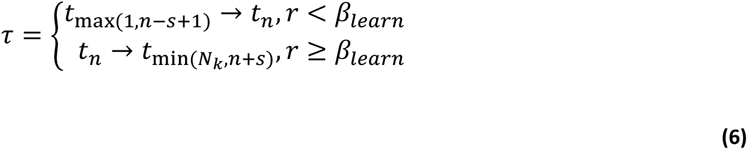
2. Update the reward estimate V over increasing indices of *τ* by computing the update equation:

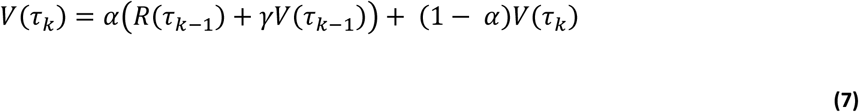 Where:

- *α* ∈ [0, 1] is the learning rate constant
- *γ* ∈ [0, 1] is the discount constant
- *R*(*t*) is the observed instantaneous reward information

This is a convex combination of the current estimate of the reward information *v*(*τ_k_*) at the next time step and the instantaneous reward information *R*(*τ_k−1_*) + *γV*(*τ_k−1_*) based on the previously observed reward signal *R*(*τ_k−1_*) and delayed previous reward estimate *γV*(*τ_k−1_*)). Equation 7 implements a form of temporal difference learning. It is sufficient to define a coarse reward signal as:

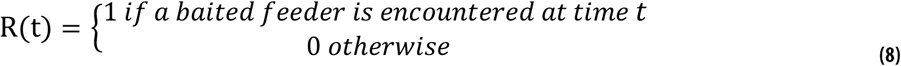

The snippet generation procedure is simply the repetition of the steps a and c of procedure (8) with *β_generate_* used instead of *β_learn_* until the sum of time subsequences durations overflows the fixed budget duration. These snippets will serve as inputs to the reservoir model of PFC described next. As illustrated in Figure 2 D and E, the replay is biased by proximity to reward, which has spatially propagated.

### Reservoir model of PFC for snippet consolidation

We model the prefrontal cortex as a recurrent reservoir network. Reservoir computing refers to a class of recurrent network models with fixed recurrent connections. The reservoir units are driven by external inputs and the network dynamics provides a high dimensional representation of the inputs from which the desired outputs can then be read out by a trained linear combination of the reservoir unit activities. The principle has been co-developed in distinct contexts as the temporal recurrent network (<u>Dominey 1995</u>), the liquid state machine (<u>Maass et al 2002</u>), and the echo state network (<u>Jaeger 2001</u>). The version that we use to model the frontal cortex employs leaky integrator neurons in the recurrent network. This model of PFC is particularly appropriate because the recurrent network generates dynamic state trajectory that will allow overlapping snippets to have overlapping state trajectories. This property will favor consolidation of a whole sequence from its snippet parts. At each time-step, the network is updated according to the following schema:

The hippocampus place-cells project into the reservoir through feed-forward synaptic connections noted *W_ffwd_*. The projection operation is a simple matrix-vector product. Hence, the input projection through feed-forward synaptic connections is defined by:

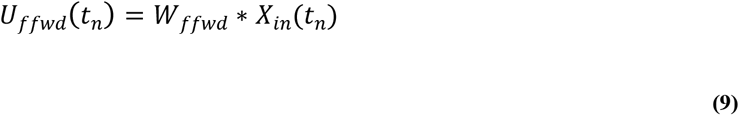

Where:

- *W_ffwd_* is a fixed connectivity matrix whose values do not depend on time.

Synaptic weights are randomly selected at the beginning of the simulation. Practically speaking (<u>Lukosevicius 2012</u>), sampling U[−1, 1] a uniform distribution is sufficient. A positive synaptic weight in a connectivity matrix models an excitatory connection and a negative weight models an inhibitory connection between two neurons (that could be implemented via an intervening inhibitory interneuron). Let N be the number of neurons in the Reservoir. Reservoir’s neurons are driven by both sensory position inputs *X_in_*(*t_n_*) and, importantly by the recurrent connections that project an image of the previous reservoir state back into the reservoir. The recurrent projection is defined as:

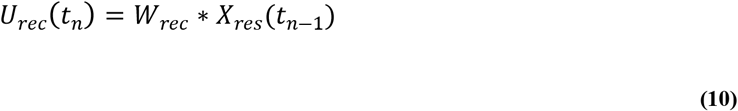

Where:

- *W_rec_* is a N by N square connectivity matrix.

Synaptic weights are drawn from a U[−1, 1] uniform distribution, scaled by a *S*(*N*; *K*) = 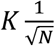 factor.

The same sign convention as in equation (9) applies for the recurrent connectivity matrix.

Self-connections (i.e. 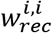 with *i* ∈ 1 … *N*) are forced to zero. *W_rec_* is also fixed and its values do not depend on time. The contributions of afferent neurons to the reservoir’s neurons is summarized by

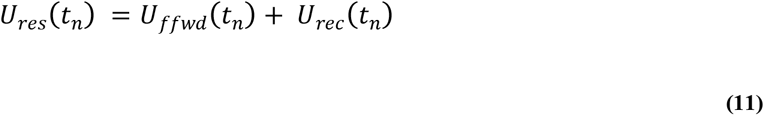

The membrane potential of the reservoir’s neurons *P_res_* then is computed by solving the following ordinary derivative equation (ODE):

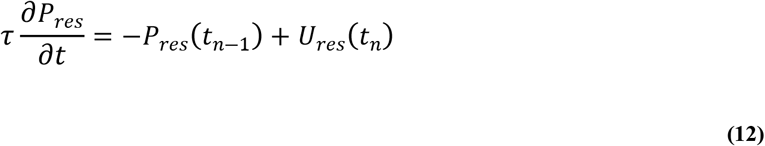

Where:

- *τ* is the neuron’s time constant. It models the resistive and capacitive properties of the neuron’s membrane.

In this article, we will consider a contiguous assembly of neurons that share the same time constant. The inverse of the time constant is called the leak rate and is noted *h*. By choosing the Euler’s forward method for solving equation(12), the membrane potential is computed recursively by the equation:

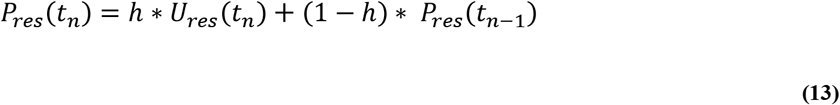

This is a convex combination between instantaneous contributions of afferents neurons *U_res_*(*t_n_*) and the previous value *P_res_*(*t_n−1_*) of the membrane potential. The current membrane potential state carries information about the previous activation values of the reservoir, provided by the recurrent weights. The influence of the history is partially controlled by the leak rate. A high leak rate will result in a responsive reservoir with a very limited temporal memory. A low leak rate will result in a slowly varying network whose activation values depend more on the global temporal structure of the input sequence.

Finally, the mean firing rate of a reservoir’s neuron is given by:

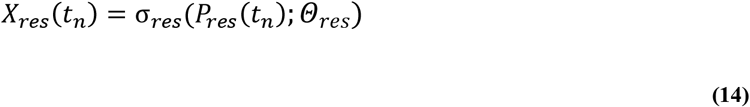

Where:

- σ*_res_* is the non-linear activation function of the reservoir neurons
- *Θ_res_* is a bias that will act as a threshold for the neuron’s activation function.

We choose a σ*_res_* ≡ *tanh* hyperbolic tangent activation function with a zero bias. Negative firing rate values represent the inhibitory/excitatory connection type in conjunction with the sign of the synaptic weight. Only the product of the mean firing rate of the afferent neuron by its associated synaptic weight is viewed by the leaky integrator neuron. See S1 High dimensional processing in the reservoir for more details on interpreting activity in the reservoir.

### Learning in Modifiable PFC Connections to Readout

Based on the rich activity patterns in the reservoir, it is possible to decode the reservoir’s state in a supervised manner in order to produce the desired output as a function of the input sequence. This decoding is provided by the readout layer and the matrix of modifiable synaptic weights linking the reservoir to the readout layer, noted *W_ro_* and represented by dash lines in Figure 3.

The readout activation pattern *X_ro_*(*t_n_*) is given by the equation:

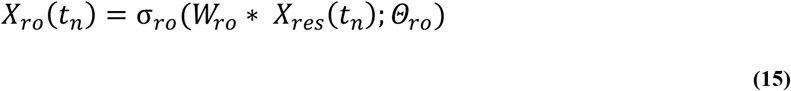

Where:

- σ*_ro_* is the non-linear activation function of the readout neurons
- *Θ_ro_* is a bias that will act as a threshold for the neuron’s activation function

We choose a σ*_ro_* ≡ *tanh* hyperbolic tangent activation function with a zero bias.

Notice that the update algorithm described above is a very particular procedure inherited from feedforward neural networks. We chose to use it because it is computationally efficient and deterministic.

Once the neural network states are updated, the readout synaptic weights are updated by using a stochastic gradient descent algorithm. By deriving the Widrow-Hoff Delta rule (<u>Widrow & Hoff 1960</u>) for hyperbolic tangent readout neurons, we have the following update equation:

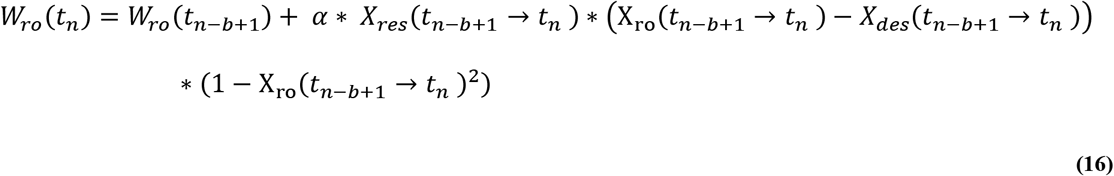

Where:

- *α* is a small positive constant called the learning rate
- *t_n_*_−*b*+1_ → *t_n_* is the concatenation of b time steps from *t_n_*_−*b*+1_to *t_n_*

When *b* = 1, equation (16) computes a stochastic gradient descent. The case when *b* > 1 is called a mini-batch gradient descent and allows one to estimate the synaptic weight gradient base on b successive observations of predicted and desired activation values. A mini batch gradient allows one to compute efficiently and robustly the synaptic weight gradient. Empirically, *b* = 32 gives satisfying results.

In this study, we will focus on the prediction of the next place-cell activation pattern:

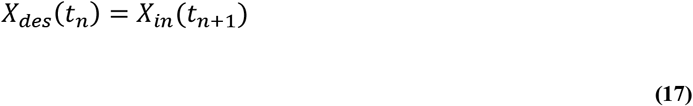

This readout is considered to take place in the striatum, as part of a cortico-striatal learning system. This is consistent with data indicating that while hippocampus codes future paths, the striatum codes actual location (<u>van der Meer et al 2010</u>).

### Training

After each trial, the model is trained using a dataset that is generated online by the snippet replay mechanism described above in the paragraph on Hippocampus replay. The readout synaptic weights are also learned online by using the learning rule described in the Learning in Modifiable PFC Connections to Readout section. The model does not receive any form of feedback from the environment and it learns place-cell activation sequences based only on random replay of snippets.

Between each sequence of the training set (snippets in our case), the states of the reservoir and readout are set to a small random uniform value centered on zero. This models a time between the replays of two snippets that is sufficiently long for inducing states in the neural network that are not correlated with the previous stimulus. This is required for having the same effect as simulating a longer time after each snippet but without having to pay the computational cost associated to this extra simulation time.

### Embodied Simulation of Sensory-Motor loop via the spatial filter

Once the model is trained, we need to evaluate its performance and the trajectories it can generate. The model is primed with the first *p* steps of the place-cell activation sequence the model is supposed to produce. This sequence is called the target sequence. Then the model’s ability to generate a place-cell activation sequence is evaluated by injecting the output prediction of the next place-cell activation pattern as the input at the next step. In this iterative procedure, the system should autonomously reproduce the trained sequence pattern of place-cell activations.

Predicted place-cell activation values might be noisy, and the reinjection of even small amounts of noise in this autonomous generation procedure can lead to divergence. We thus employ a procedure that determines the location coded by the place-cell activation vector output, and reconstructs a proper place-cell activation vector coding this location. We call this denoising procedure the spatial filter as referred to in Figure 3.

We model the rat action as ‘reaching the most probable nearby location’. Since only the prediction of the next place-cell activation pattern *η* is available, we need to estimate the most probable point *s^∗^*(*t_n_*_+1_) = (*x^∗^*(*t_n_*_+1_), *y^∗^*(*t_n_*_+1_)). From a Bayesian point of view, we need to determine the most probable next location *s*(*t_n_*_+1_), given the current location *s*(*t_n_*) and the predicted place-cell activation pattern *η*(*t_n_*). We can state our problem as:

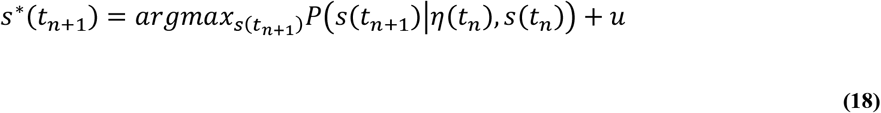

Where:

- *u* is a noise function sampling a uniform distribution *𝒰*(0, *m*)

*u* is useful at least in degenerate cases when a zero place-cell activation prediction generates an invalid location coding. It is also used for biasing the generation procedure and to explore other branches of the possible trajectories the model can generate as described in section Evaluating Behavior with Random walk.

The system is then moved to this new location *s^∗^* and a new noise/interference free place-cell activation pattern is generated by the place-field model. We refer to this place-cell prediction/de-noising method as the spatial filter, which emulates a sensory-motor loop for the navigating rat in this study. Figure 3 depicts this sensory motor loop.

### Evaluating Behavior with Random walk

Once the model has been trained, it is then primed with place-cell activation inputs corresponding to the first few steps of the trajectory to be generated. The readout from the PFC reservoir generates the next place-cell activation pattern in the trajectory, which is then reinjected into the reservoir via the spatial filter, in a closed loop process. This loop evaluation procedure is called *autonomous* generation. In order to evaluate the model in a particular experimental condition, several instances of the same model are evaluated multiple times in a random walk procedure. The batch of generated trajectories (typically 1000) are accumulated in a stencil buffer which acts as a two dimensional histogram showing the most frequently generated trajectories. The arena is drawn with its feeders and a vector field is computed from trajectories in order to show the main direction of these trajectories. Trajectories are superimposed and summed, resulting in a two-dimensional histogram representing the space occupied by trajectories. Figure 4 shows an example of random walk trajectories, illustrating the model’s ability to autonomously generate a long and complex sequence when learning without snippets.

**Figure 4:**
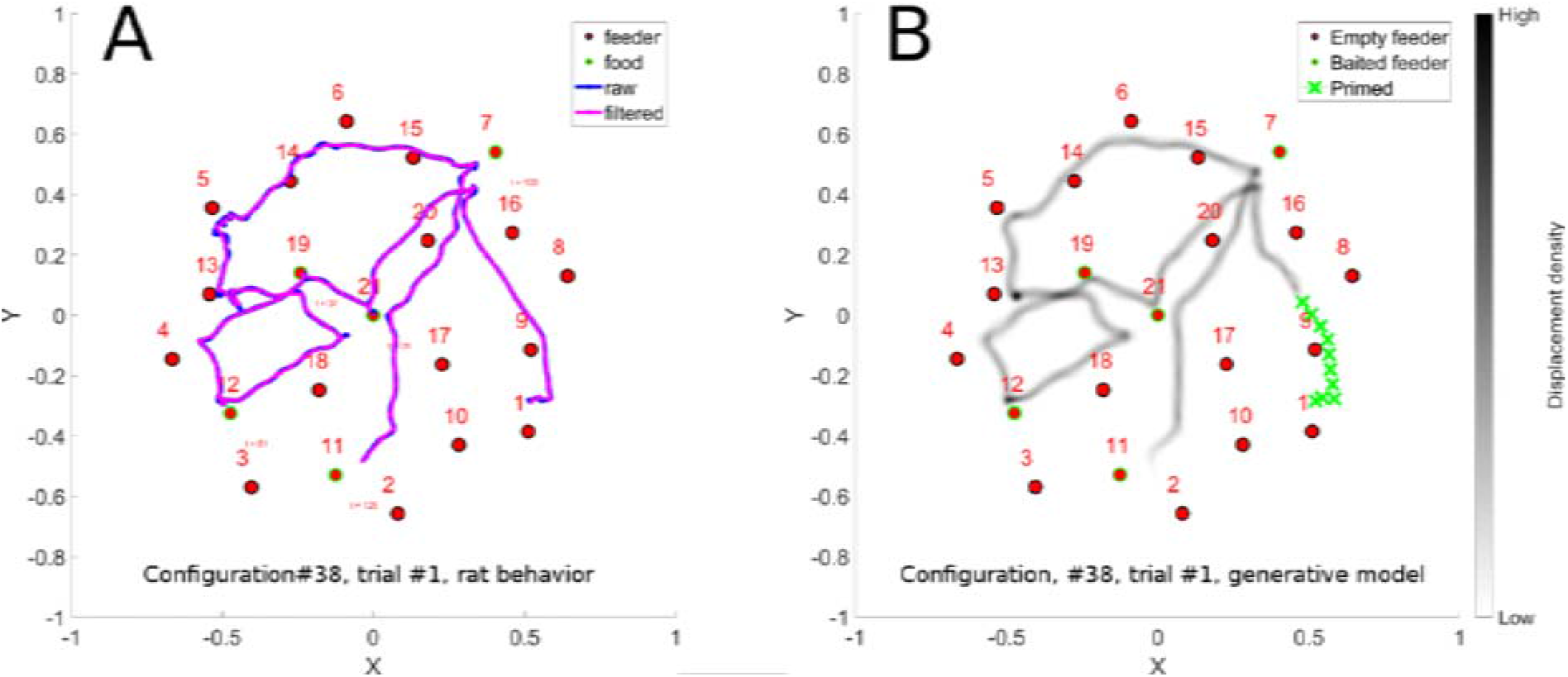
Sequence learning. Panel A illustrates a long convoluted trajectory taken by a rat in configuration 38. Panel B illustrates the probability maps of trajectories generated by the trained model in autonomous sequence generation mode. Note that there are two locations where the trajectory crosses itself, which introduces ambiguity that the model is able to disambiguate. This illustrates that the model is well able to learn such sequences.

In cases where small errors in the readout are reinjected as input, they can be amplified, causing the trajectory to diverge. It is possible to overcome this difficulty by providing as input the expected position at each time step instead of the predicted position. The error/distance measurement can still be made, and will quantify the diverging prediction, while allowing the trajectory generation to continue. This method is called *non-autonomous* generation and it evaluates only the ability of a model to predict the next place-cell activation pattern, given an input sequence of place-cell activations.

### Comparing produced and ideal sequences using Discrete Fréchet distance

The joint PFC-HIPP model can be evaluated by comparing an expected place-cell firing pattern with its prediction by the readout layer. At each time step, an error metric is computed and then averaged over the duration of the expected neurons firing rate sequence. The simplest measure is the mean square error. This is the error that the learning rule described in equation (16) minimizes.

Although the model output is place-cell coding, what is of interest is the corresponding spatial trajectory. A useful measurement in the context of comparing spatial trajectories is the discrete Fréchet distance. It is a measure of similarity between two curves that takes into account the location and ordering of the points along the curve. We use the discrete Fréchet distance applied to polygonal curves as initially described in <u>Eiter and Mannila (1994)</u>. In (<u>Wylie 2013</u>) the Discrete Fréchet distance *F* between two curves *A* and *B* is defined by:

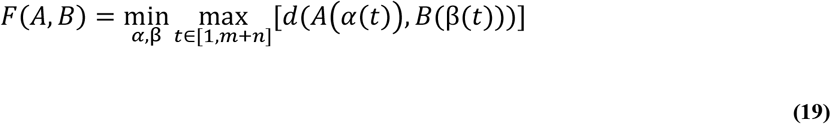

Where *d*(.,.) is the Euclidean distance, *m* is the number of steps of the curve A, *n* is the number of steps of the curve B, and *α*, *β* are reparametrizations of the curves A and B. Parameterization of this measure is described in more detail in S1 Frechet distance parameters.

## Results

For robustness purposes, results are based on a population of neural networks rather than a single instance. The population size is usually 1000 for evaluating a condition and the metrics described above are aggregated by computing their mean *μ*(.) and standard-deviation *σ*(.). For convenience, we define a custom score function associated to a batch of coherent measurements as:

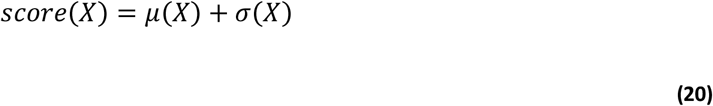

Results having a low mean and standard deviation will be reported as low score whilst other possible configurations will result in a higher score. We choose this method rather than Z-score, which penalizes low standard deviations. We first established that the model displays standard sequence learning capabilities (e.g. illustrated in Figure 4) and studied parameter sensitivity (see S1 Basic Sequence learning and parameter search), and then addressed consolidation from replay.

### Consolidation from snippet replay

The model is able to learn and generate navigation sequences from place-cell activation patterns. The important questions is whether a sequence can be learned by the same model when it is trained on randomly presented snippets, instead of the continuous sequence.

In this experiment, no reward is used, and thus each snippet has equal chance of being replayed. The only free parameter is the snippet size. In order to analyze the reservoir response, we collect the state-trajectories of reservoir neurons when exposed to snippets. Recall that the internal state of the reservoir is driven by the external inputs, and by recurrent internal dynamics, thus the reservoir adopts a dynamical state-trajectory when presented with an input sequence. Such a trajectory is visualized in Figure 5D. This is a 2D (low dimensional) visualization, via PCA, of the high dimensional state transitions realized by the 1000 neurons reservoir as the input sequence corresponding to ABCDE is presented. Panels A-C illustrate the trajectories that the reservoir state traverses as it is exposed to an increasing number of randomly selected snippets generated for the same ABCDE sequence. We observe that as snippets are presented, the corresponding reservoir state-trajectories start roughly from the same point because of the random initial state of the reservoir before each snippet is replayed. Then the trajectories evolve and partially overlap with the state-trajectory produced by the complete sequence. In other words, snippets quickly drive the reservoir state from an initial random activation (corresponding to the grey area at the center of each panel) onto their corresponding locations in the reservoir activation state-trajectory of the complete sequence. Replaying snippets at random thus has no negative impact because the reservoir states overlap when snippet trajectories overlap.

**Figure 5:**
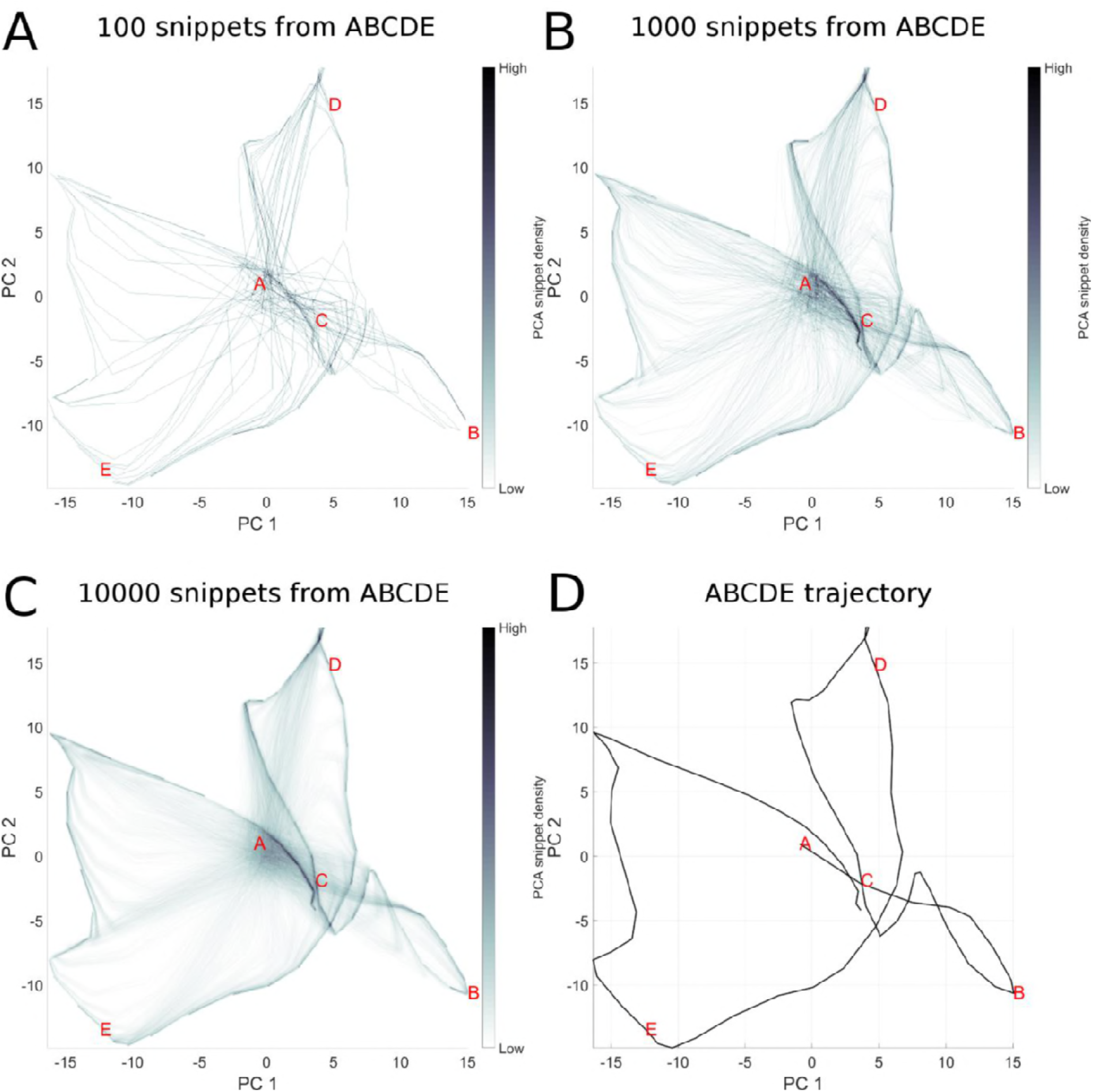
Illustration of snippet integration in reservoir state space. Here we visualize the high dimensional reservoir space in a low (2D) PCA space, in order to see how pieces (snippets) of the overall sequence are consolidated. In this experiment, the sequence ABCDE is broken into snippets, which are then used to train the model. The challenge is that only local structure is presented to the model, which must consolidate the global structure. Panels A-C represent the state trajectory of reservoir activation after 100, 1000 and 1000 snippets. While each snippet represents part of the actual trajectory, each is taken out of its overall spatial context in the sequence. Panel D represents the trajectory of reservoir state during the complete presentation of the intact sequence. Panel C reproduces this trajectory, but in addition we see “ghost” trajectories leading to the ABCDE trajectory. These ghost elements represent the reservoir state transitions from an initial random state as the first few elements of each snippet take the reservoir from the initial undefined state onto the component of the ABCDE trajectory coded by that snippet.

Thus, we see that the state trajectories traversed by driving the reservoir with snippets overlaps that from the original intact sequence. See further details of sequence learning by snippet replay in S1 Sequence complexity effects on consolidation.

### Longer paths are rejected

Here we examine how using reward proximity to modulate snippet replay probability distributions (as described in the hippocampal replay description) allows the rejection of longer, inefficient paths between rewarded targets. In this experiment, 1000 copies of the model are run 10 times. Each is exposed to the reward modulated replay of two sequences ABC and ABD having a common prefix AB as illustrated in Figure 6. The model is exposed to a random replay of sequences ABC and ABE. The random replay is not uniform and takes into account the reward associated with a baited feeder when food was consumed. Snippets close to a reward have more chance to be replayed and thus to be consolidated into a trajectory.

**Figure 6:**
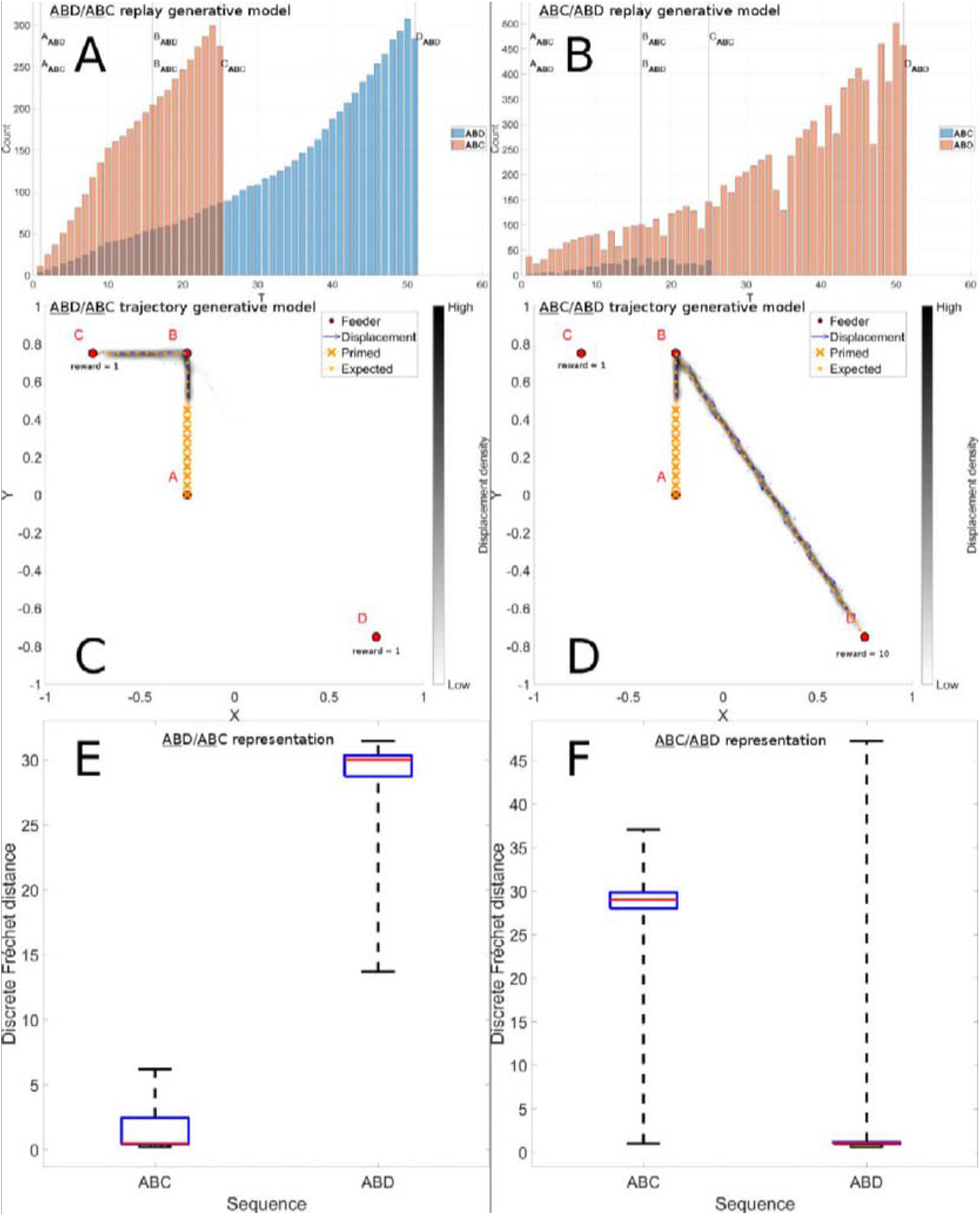
Longer paths are rejected. Panels A and B illustrate snippet counts for T maze trajectories pictured in panels C and D. In Panel C, sequences begin at location A, and rewards are given at locations C and D. Based on the reward proximity and propagation, there is a higher probability of snippets being selected along path AC than path AD. This is revealed in panel A, a histogram of snippets for the sequences ABC (in Blue) and ABD (in Orange). Panels B and D illustrate how distance and reward intensity interact. By increasing the strength of the reward, a longer trajectory can be rendered virtually shorter and more favored, by increasing the probability that snippets will be selected from this trajectory, as revealed in Panel B. Panel E and F confirm a neat tendency to generate autonomously sequences significantly similar to the ABC and ABD sequence respectively (p-value = 0).

Panel A in Figure 6 illustrates the distribution of snippets selected from the two sequences, ABC in pink and ABD in blue. At the crucial point of choice at location B, the distribution of snippets for sequence ABC largely outnumbers those for sequence ABD. This is due to the propagation of rewards respectively from points C and D. Per design, rewards propagated from a more proximal location will have a greater influence on snippet generation. Panel C shows the 2D histogram of autonomously generated sequences when the model is primed with the initial sequence prefix starting at point A. We observe a complete preference for the shorter sequence ABC illustrated in panel E.

The snippet generation model described above takes into account the location of rewards, and the magnitude of rewards. Panel B illustrates the distribution of snippets allocated to paths ABC and ABD when a 10x stronger reward is presented at location D. This strong reward dominates the snippet generation and produces a distribution that strongly favors the trajectory towards location D, despite its farther distance. Panel F illustrates the error mesures for model reconstruction of the two sequences and confirms this observation. This suggests an interesting interaction between distance and reward magnitude. For both conditions, distances to the expected sequence have been measured for every trajectory generated (10 000 for ABC and 10 000 for ABD). Then a Kruskall Wallis test confirms (p-value ~= 0) for both cases that trajectories generated autonomously are significantly more accurate for the expected trajectories (i.e. ABC when rewards are equal and ABD when reward at D is x10).

### Novel efficient sequence creation

Based on the previously demonstrated dynamic properties, we determined that when rewards of equal magnitudes are used, the model would favor shorter trajectories between rewards. We will now test the model’s ability to exploit this capability, in order to generate a novel and efficient trajectory from trajectories that contain sub-paths of the efficient trajectory. That is, we determine whether the model can assemble the efficient subsequences together, and reject the longer inefficient subsequences in order to generate a globally efficient trajectory. Figure 1 (Panel A) illustrates the desired trajectory that should be created without direct experience, after experience with the three trajectories in panels B-D that each contain part of the optimal trajectory (red), which will be used to train the model.

**Figure 7:**
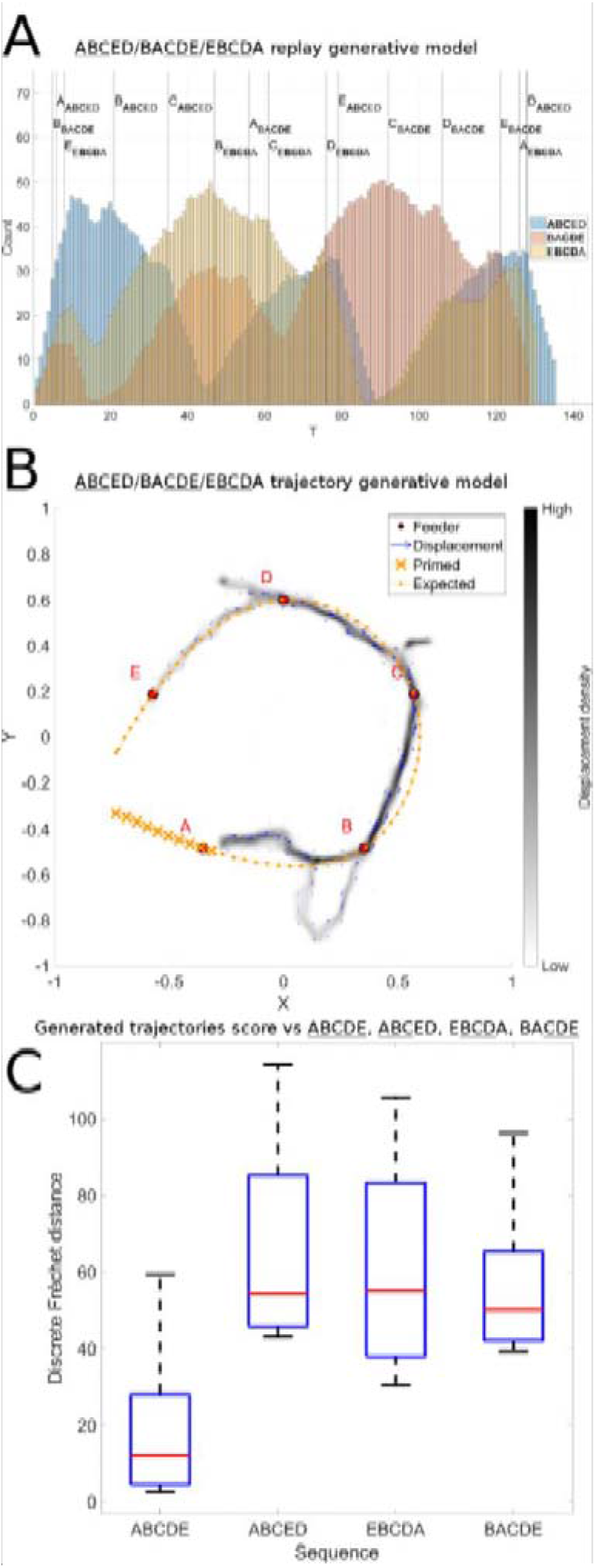
Efficient Sequence Synthesis. A. Distribution of snippets drawn from the sequences illustrated in Figure 1 B, C and D. Globally we observe snippet selection favors snippets from the beginning of sequence ABCED (blue), the middle of EBCDA (yellow), and the end of sequence BACDE (pink), which corresponds exactly to the efficient subsequences (ABC, BCD, and CDE) of these three sequences. This distribution of snippets is used to train the model. The results of the training are illustrated in panel B. Here we see a 2D histogram of sequences generated by the model in the ABCDE recombination experiment. 10 batches of 100 reservoirs each were trained and each model instance was evaluated 10 times with noise. Panel C confirms that the trajectories generated autonomously are significantly more similar to the target sequence ABCDE (p-value = 5.9605e-08) These results are very robust and satisfying, demonstrating that our hypothesis for efficient sequence discovery based on reward-modulated replay is validated.

The reward-biased replay is based on the following trajectories: (1) <u>ABC</u>ED that contains the ABC part of the ABCDE target sequence, (2) E<u>BCD</u>A that contains the BCD part of the ABCDE target sequence, and (3) BA<u>CDE</u> that contains the CDE part of the ABCDE target sequence. Figure 7 A illustrates how the hippocampal replay model generates distributions of snippets that significantly favor the representation of the efficient subsequences of each of the three training sequences. This is revealed as the three successive peaks of snippet distributions on the time histogram for the blue (<u>ABC</u>ED) sequence, favoring its initial part ABC, the yellow (E<u>BCD</u>A) sequence, favoring its middle part BCD, and the pink (BA<u>CDE</u>) sequence, favoring its final part CDE. When observing each of the three color-coded snippet distributions corresponding to each of the three sequences we see that each sequence is favored (with high replay density) precisely where it is most efficient. Thus, based on this distribution of snippets that is biased towards the efficient subsequences, the reservoir should be able to extract the efficient sequence.

This is shown in panel B, which illustrates the autonomously generated sequences for 1000 instances of the model executed 10 times each. The spatial histogram reveals that the model is able to extract and concatenate the efficient subsequences to create the optimal path, though it was never seen in its entirely in the input. Panel C illustrate the significant differences in performance between the favored efficient sequence vs. the three that contain non-efficient subsequences. A Kruskal-Wallis test confirms these significant differences reconstruction error for the efficient vs non-efficient sequences (maximum p = 5.9605e-08).

### Reverse replay

In (<u>Carr et al 2011</u>), hippocampus replay during SWR is characterized by the activation order of the place-cells which occurs in backward and forward direction. We hypothesize that reverse replay allows the rat to explore a trajectory in one direction but consolidate it in both directions. This means that an actual trajectory, and its unexplored reverse version, can equally contribute to new behavior. Thus fewer actual trajectories are required for gathering information for solving the TSP problem. A systematic treatment of this effect on learning can be seen in S1 Analysis of different degrees of reverse replay.

We now investigate how reverse replay can be exploited in a recombination task where some sequences are experienced in the forward direction, and others in the reverse direction, with respect to the order of the sequence to be generated. We use the same setup as described above for novel sequence generation, but we invert the direction of sequence EBCDA in the training set. Without EBCDA, the model is not exposed to sub trajectories linking feeders B to C and C to D and the recombination cannot occur. We then introduce a partial reverse replay, which allows snippets to be played in forward and reverse order. This allows the reservoir to access segments BC and CD (even though they are not present in the forward version of the experienced trajectory.

Figure 8 illustrates the histogram of sequence performance for 10000 runs of the model (1000 models run 10 times each) on this novel sequence generation task with and without 50% reverse replay. We observe a significant shift towards reduced errors (i.e. towards the left) in the presence of reverse replay.

**Figure 8:**
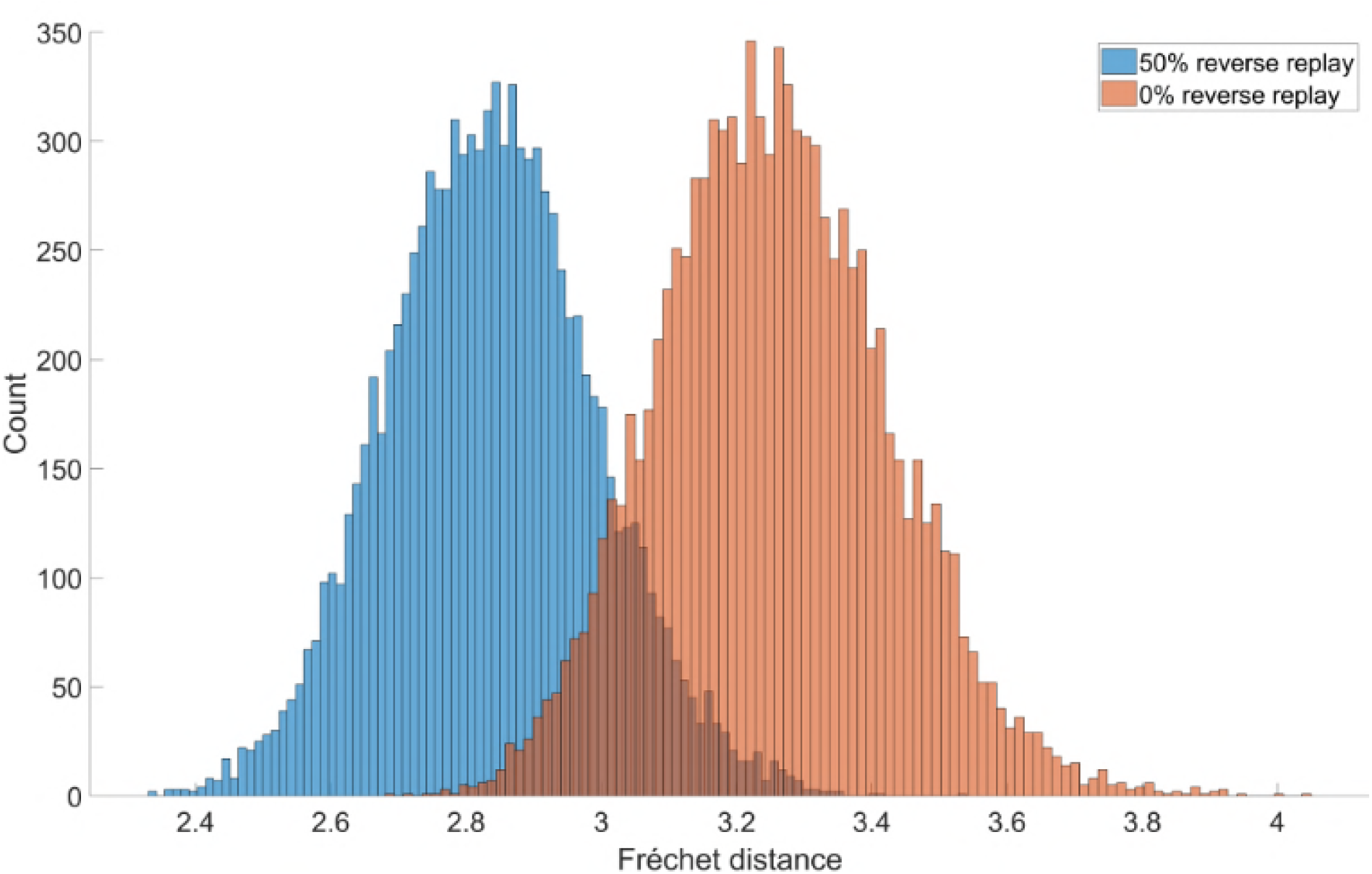
Reverse replay facilitates efficient sequence discovery. Using the same sequences illustrated in Figure 1, we reversed the direction of sequence EBCDA, and then tested the model’s ability to synthesize the ABCDE sequence from <u>ABC</u>ED A<u>CDB</u>E and BA<u>CDE</u>

We then examine a more realistic situation based on the observation of spontaneous creation of “shortcuts” described in (<u>Gupta et al 2010</u>). The model is exposed to a random replay of snippets extracted from two trajectories having different direction (clockwise CW and counter clockwise CCW). The system thus experiences different parts of the maze in different directions. We examine whether the use of reverse replay can allow the system to generate novel shortcuts.

The left and right trajectories used for training are illustrated in Figure 9A and B. In A, the system starts at MS, head up and to the left at T2 (counter clockwise) and terminates back at MS. In B, up and to the right (clockwise) again terminating at MS. Possible shortcuts can take place at the end of a trajectory at MS as the system continues on to complete the whole outer circuit rather than stopping at MS. We can also test for shortcuts that traverse the top part of the maze by starting at MS and heading left or right and following the outer circuit in the CW or CCW direction, thus yielding 4 possible shortcuts. The model is trained with snippets from the sequences in A and B using different random replay rates, and evaluated in non-autonomous mode with sequences representing the 4 possible types of shortcut. Figure 16 C shows with no reverse replay, when attempting the CCW path, there is low error until the system enters the zone that has only been learned in the CW direction. In panel D, with 50% reverse replay, this error is reduced and the system can perform the shortcut without having experienced the right hand part in the correct direction. Thus, in the right hand part of the maze, it is as if the system had experienced this already in the CCW direction, though in reality this has never occurred, but is simulated by the reverse replay. This illustrates the utility of mixed forward and reverse replay. Panel E illustrates the difficulty when 100% reverse replay is used. Figure 16F illustrates the reconstruction errors for a shortcut path as a function of degree of reverse The trajectory is evaluated in non-autonomous mode and the position of the agent necessarily follows the target trajectory. In this case, the expected trajectory describes a CCW path. Results are not significantly different with a replay rate 25% and 75% (p = 0.02), where the best performance is observed, and all the other conditions are significantly different (p ≤ 1.1921e-07). This phenomenon was obtained for the 4 possible shortcuts.

**Figure 9:**
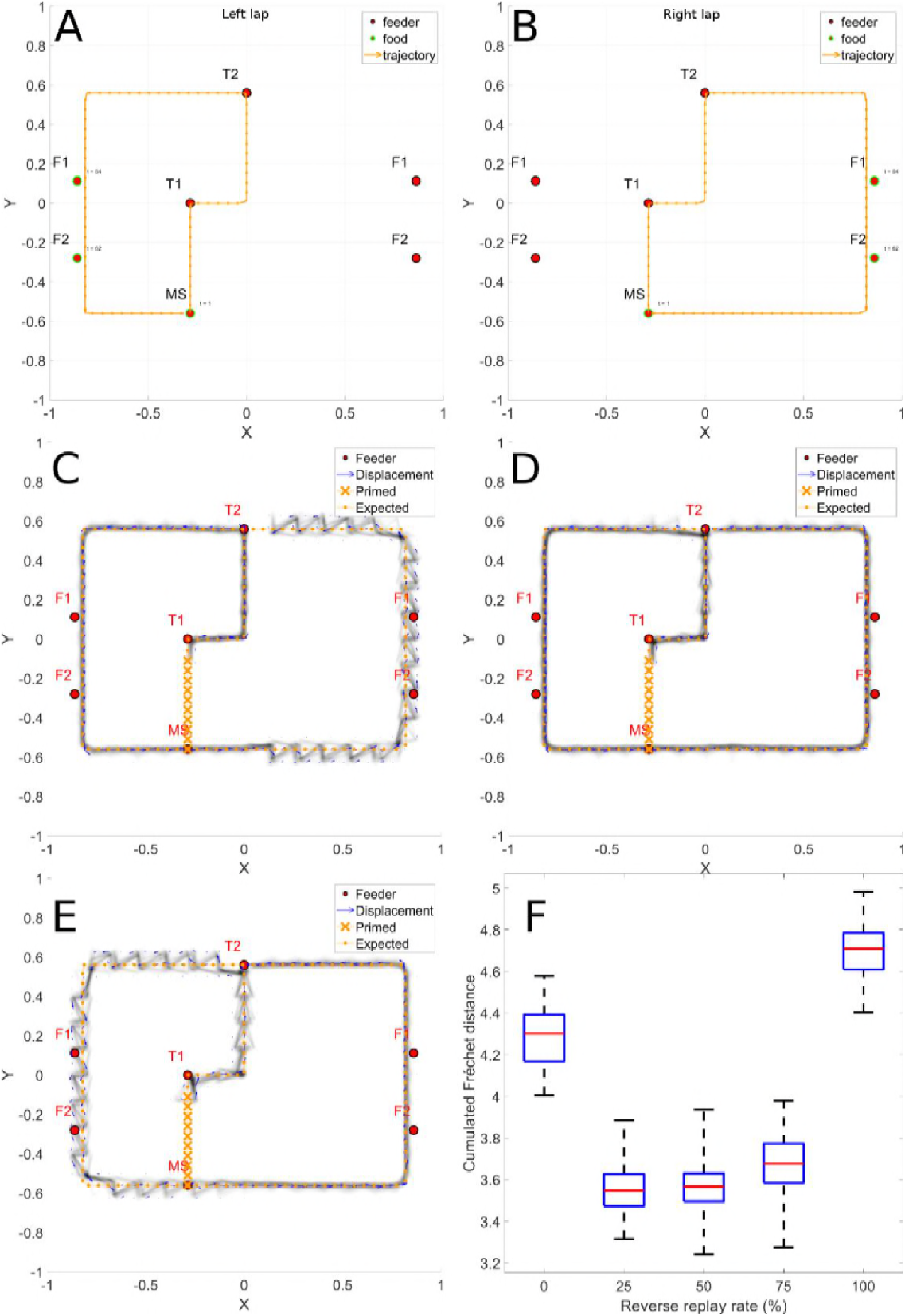
Reverse replay allows novel shortcut path generation. Panels A and B illustrate the trajectories for left and right trajectories, based on Gupta et al. After training on these two trajectories, we test the ability to generate a shortcut that makes the complete outer loop in one direction. Panel C – without reverse replay, significant spatial errors are revealed when the system attempts to complete the counter-clockwise loop on the right side of the maze Panel D illustrates the beneficial effects of reverse replay during trajectory learning. Panel E illustrates the effect of a model training with 100% reverse replay. It is similar to using a 0% reverse replay but the effect is observed on the left lap trajectory part. Panel F – when reverse replay is introduced, this error is attenuated.

### Effects of Consolidation and Reverse Replay

The model demonstrates the ability to accumulate and consolidate paths over multiple trials, and to exploit reverse replay. Here we examine these effects on the more extensive and variable dataset extracted from rat behavior (<u>de Jong et al 2011</u>). We show the positive effects of replay on trajectories from rats trying to optimize spatial navigation in the TSP task. In the prototypical TSP behavior, in a given configuration of baited wells, on successive trials the rat traverses different efficient subsequences of the overall efficient sequence, and then finally puts it all together and generates the efficient sequence. This suggests that as partial data about the efficient sequence are successively accumulated, the system performance will successively improve. To explore this, the model is trained on navigation trajectories that were generated by rats in the TSP task. We selected data from configurations where the rats found the optimal path after first traversing subsequences of that path in previous trials. Interestingly, these data contain examples where the previous informative trails include traversal of part of the optimal sequence in either the forward or reverse directions, and sometimes both (see S Rat navigation data). We trained training the model with random replay of combinations of informative trials where informative trials are successively added, in order to evaluate the ability of the model to successively accumulate information. For each combination of informative trials, the random replay is evaluated with 0%, 25%, 50%, 75% and 100% of reverse replay rate in order to assess the joint effect of random replay and combination of informative trials. The model is then evaluated in non-autonomous mode with the target sequences that consist in a set of trajectories linking the baited feeders in the correct order. An idealized sequence is added to the target sequence set because trajectories generated by the rat might contain edges that do not relate the shortest distance between two vertices. Agent’s moves are restricted to a circle having a 10 cm radius.

Figure 10 illustrates the combined effects of successive integration of experience and its contribution to reducing error, and of the presence of different mixtures of forward and reverse replay. The ANOVA revealed that there is a significant effect for combination (F(2, 585) = 32.84, p < 0.01), as performance increases with exposure to more previous experience (Panel A). There is also a significant effect for reverse replay rate (F(4, 585) = 3.71, p ≤ 0.01), illustrated in Panel B. There was no significant interaction between consolidation and replay direction (F(8, 585) = 0.03, p = 1). This indicates that when trained on trajectories produced by behaving rats, the model displays the expected behavior of improving with more experience, and of benefitting from a mixture of forward and reverse replay.

**Figure 10:**
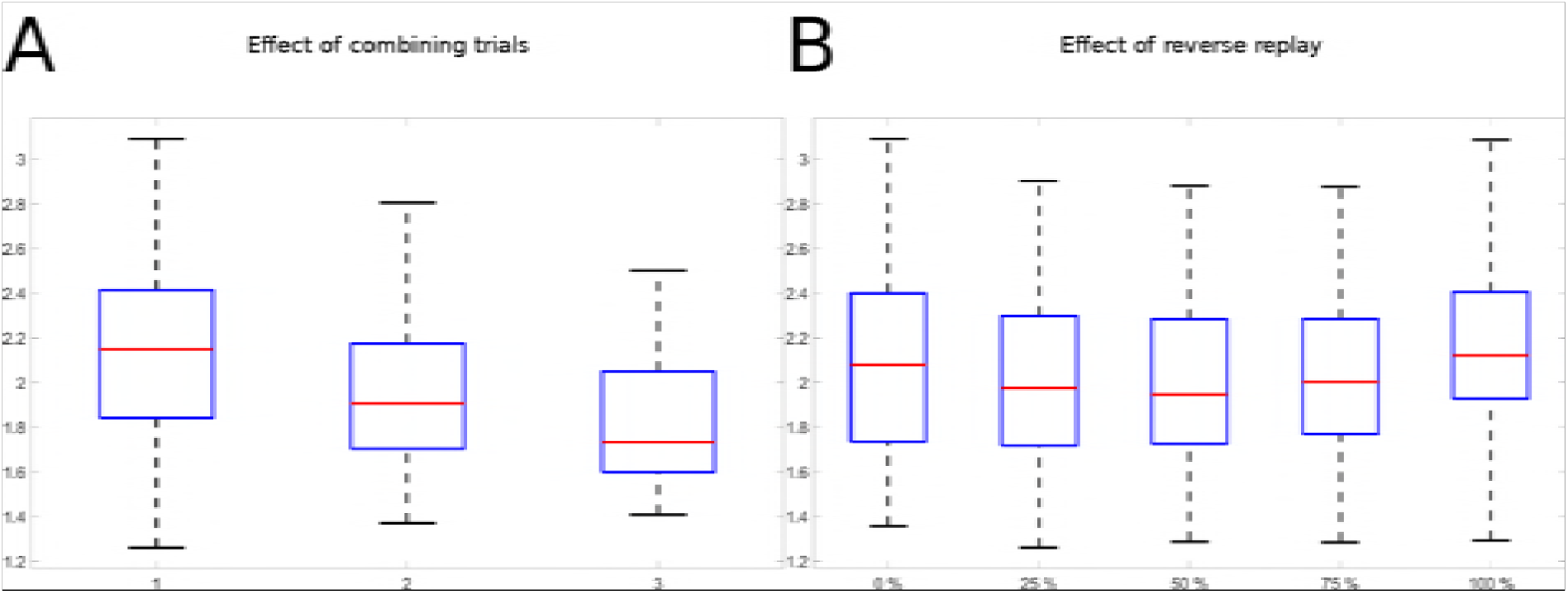
Consolidation and reverse replay applied to behavioral data. Measured variable is Frechet distance between generated and desired sequence. Data from the rat TSP configurations are used for training and testing the model. A. Effects of consolidation: as successive trials are added to the replay repertoire; the trajectory reconstruction error is significantly reduced. B. Effects of reverse replay: as reverse replay is introduced in snippet formation for training the PFC model, reconstruction error is significantly reduced.

## Discussion

We tested the hypothesis that hippocampus replay observed during sharp wave ripple events in the awake animal can play a role in memory consolidation by exposing the prefrontal cortex between successive trials to short subsequences of place-cell activation patterns. This replay can potentially play a crucial role in learning, essentially by generating synthetic data (based on experience) for training the system. The behavior of interest is a form of spatial navigation trajectory optimization in a task, mimicking the well-known traveling salesperson problem. It is a NP-Hard problem and finding an exact solution would require significant time and computing resources. Nevertheless, it has been observed that a rat was able to quickly solve simplified versions of this problem (<u>Bureš et al 1992</u>, <u>de Jong et al 2011</u>). The idea of exploiting replay in navigation sequence learning has been demonstrated to have a positive influence on learning (<u>Johnson & Redish 2005</u>), and here we go beyond this by further exploiting reward structure in the replay.

In the behavior of interest, rats are observed to converge quickly to a near-optimal path linking 5 baited food wells in a 151cm radius open arena. During their successive approximation to the optimal path, the rats often traversed segments of the optimal trajectory, as well as non-optimal segments. Observing this behavior, we conjectured the existence of neural mechanisms that would allow the optimal segments to be reinforced and the non-optical segments to be rejected, thus leading to the production of the overall near-optimal trajectory. The overall mechanism we propose can be decomposed into two distinct neural systems. The first is a replay mechanism that favors the representation of snippets that occurred on these optimal segments, and that in contrast will give reduced representation to snippets that correspond to non-optimal trajectory segments. Here we demonstrate a simple but powerful method based on spatial reward propagation that implements this mechanism. Interestingly, this characterization of replay is broadly consistent with the effects of reward on replay observed in behaving animals (<u>Ambrose et al 2016</u>).

The second neural system required to achieve this integrative performance is a sequence learning system that can integrate multiple subsequences (i.e. snippets) into a consolidated representation, taking into consideration the probability distributions of replay so as to favor more frequently replayed snippets. Here we considered a well-characterized model of sequence learning based on recurrent connections in prefrontal cortex that is perfectly suited to meet the sequence learning requirements.

### Replay mechanism

Replay is modeled using a procedure that randomly selects a subset of place-cells coding part of a sequence, and outputs this snippet while taking into account the proximity of this snippet to a future reward. Each time a reward is encountered, it is taken into consideration in generating the snippet, and reward value is propagated backwards along the sequence, thus implementing a form of spatio-temporal credit assignment. This can be viewed in the figures 2, 6 and 7 illustrating the snippet probability densities. The replay mechanism also implements a second feature observed in animal data, which is a tendency to replay snippets in reverse order. These two features of the replay model correspond to what is observed in the rat neurophysiology, and they also make fundamental contributions to the model’s ability to converge on an efficient navigation path. This extends previous demonstrations of the value of replay to include reward-modulated optimization (<u>Johnson & Redish 2005</u>).

### Reservoir network

Reservoir computing exploits the spatio-temporal dynamics of recurrently connected neurons that are sensitive to the spaiotemporal structure of input sequences (<u>Dominey 1995</u>, <u>Jaeger & Haas 2004</u>, <u>Maass et al 2002)</u>. The frontal cortex has been demonstrated to operate on these reservoir properties (<u>Enel et al 2016</u>). Here we demonstrated how a reservoir model of PFC meets two requirements for sequence learning: First, it can concatenate randomly replayed subsequences (snippets) in order to generate the compete original sequence. Second, it is sensitive to the statistics of replay, and thus can learn to ignore rare snippets (which correspond to snippets on inefficient subsequences, far from rewards) thus learning to optimize.

### Effects of reward

The instantaneous reward information acquired during a past experience is used for recursively updating the snippet replay likelihood in the hippocampus model. The resulting time distribution of snippets features multiple modes defined around the moments that rewards are obtained. This creates a reward gradient and allows sensory-motor associations to be learned by the prefrontal cortex and striatum model. This is a novel form of reinforcement learning, and the main effect of reward combined with hippocampus replay is to reinforce existing efficient paths between rewards. A secondary effect of rewards could be observed when rewards are sufficiently close for allowing a mutual contribution to the snippet replay likelihood surrounding the moments associated with reward delivery. Thus, we predict that a cluster of reward sites will have the effect of propagating the reward information farther than a single reward.

### Effects of reverse replay

The reverse replay mechanism has a dual effect. First, a snippet replayed in reverse order will play the role of a short-term support for the reward propagation in reverse direction in the snippet replay likelihood online learning. Place-cell activation sequences leading to a nearby reward are represented more frequently and earlier than other paths, and allow less efficient sub-paths to be rejected. It results in a rewarding path selection mechanism by implementing a form of saliency based on reward information. This is a form of spatio-temporal credit assignment that allows to take advantage of the reservoir network ability to combine multiple snippets into a whole sequence. We showed that it is possible to consolidate multiple sequences featuring parts of the same underlying optimal sequence into one efficient sequence and to generate it autonomously. Second, when the snippet replay likelihood is learned, a non-zero reverse replay rate allows the prefrontal cortex to be exposed to sequences of place-cell activations in both forward and reverse direction. This results in sequence learning in both directions while having experienced a place-cell activation sequence in one direction only. These results can be tested experimentally by recording place cells activities in SWR during the task.

### Conclusions and limitations

The model we studied here is able to mimic the rat’s ability to find good approximations to the traveling salesperson problem by taking advantage of recent rewarding experiences for updating a trajectory generative model using hippocampus awake replay. We showed that reverse replay allows the agent to reduce the TSP task complexity by considering an undirected graph where feeders are vertices and trajectories are the edges instead of a directed graph. In this case, autonomous sequence generation is no longer possible but the information available in each prediction of the prefrontal cortex contains the expected locations. This allows the building of a navigation policy taking into account the salient actions suggested by the prefrontal cortex predictions, which are learned from hippocampus replay.

## Acknowledgements

Work supported by NSF-ANR CRCNS (#1429929, Spaquence)

